# SpaCell: integrating tissue morphology and spatial gene expression to predict disease cells

**DOI:** 10.1101/837211

**Authors:** Xiao Tan, Andrew Su, Minh Tran, Quan Nguyen

**Affiliations:** Institute for Molecular Bioscience, The University of Queensland, Brisbane, 4072, Australia

## Abstract

**Motivation:** Spatial transcriptomics technology is increasingly being applied because it enables the measurement of spatial gene expression in an intact tissue along with imaging morphology of the same tissue. However, current analysis methods for spatial transcriptomics data do not use image pixel information, thus missing the quantitative links between gene expression and tissue morphology.

**Results:** We developed an user-friendly deep learning software, SpaCell, to integrate millions of pixel intensity values with thousands of gene expression measurements from spatially-barcoded spots in a tissue. We show the integration approach outperforms the use of gene count alone or imaging data alone to create deep learning models to identify cell types or predict labels of tissue images with high resolution and accuracy.

**Availability:** The SpaCell package is open source under a MIT license and it is available at https://github.com/BiomedicalMachineLearning/SpaCell

## 1 Introduction

Spatial transcriptomics (ST) technology is emerging as an important platform for measuring molecular biological processes at the tissue level (Burgess, 2019). Different from other genomics technologies, ST does not require dissociating cells from the original tissue. Molecular measurements can be mapped back to the spatial location of the cells in tissue via spatial barcodes, adding a novel spatial data dimension to gene expression data. Moreover, platforms such as Slide-seq generate a tissue image and a gene expression profile of the same tissue, allowing the integration of tissue morphology and spatial gene expression (**?**).

However, incorporating imaging data to gene expression data is a new analysis area, while current analysis pipelines mainly focus on using expression values but not image pixel values. Image pixel intensity data contain informative features that can be used for diagnosing diseases such as for cancer staging (Coudray *et al.*, 2018). Although machine learning methods exist for analysing imaging data (Komura and Ishikawa, 2018), these methods do not utilise molecular data. Advances in genomics technologies create new data types for novel machine learning applications to combine molecular measurements with image pixel data to characterise tissue morphological images beyond pathological annotation (Hekler *et al.*, 2019). Existing methods for spatial data analysis, however, use gene expression, but not image pixel information (Navarro *et al.*, 2017; Dries *et al.*, 2019). We developed SpaCell with a comprehensive workflow to utilise both pixel and gene expression data to train neural network (NN) models for cell-type and disease-stage classification.

## 2 Main workflow

SpaCell’s workflow (Fig. 1) starts with two-stream data preprocessing. For image preprocessing, SpaCell first removes any colour cast, which is the background difference from the white background in the H&E image, then performs stain normalisation to overcome inconsistencies in the staining process (Macenko *et al.*, 2009) (Supp. Methods). Then, high images are tiled into small tiles and the tiles are resized to 299 × 299 pixels, where each tile contains one spot. To increase model performance and generalisability, SpaCell performs random rotation and Z-transform of the tiled images for each training step. For count matrix preprocessing, gene counts are mapped read counts to each spatial transcriptomics spot, recovered by spatial barcodes. A large range of programs developed for single-cell data analysis are available for users to process and normalise count data. SpaCell has built-in and fast options to remove unreliably detected spots and genes, followed by library-size normalisation.

**Fig 1.**
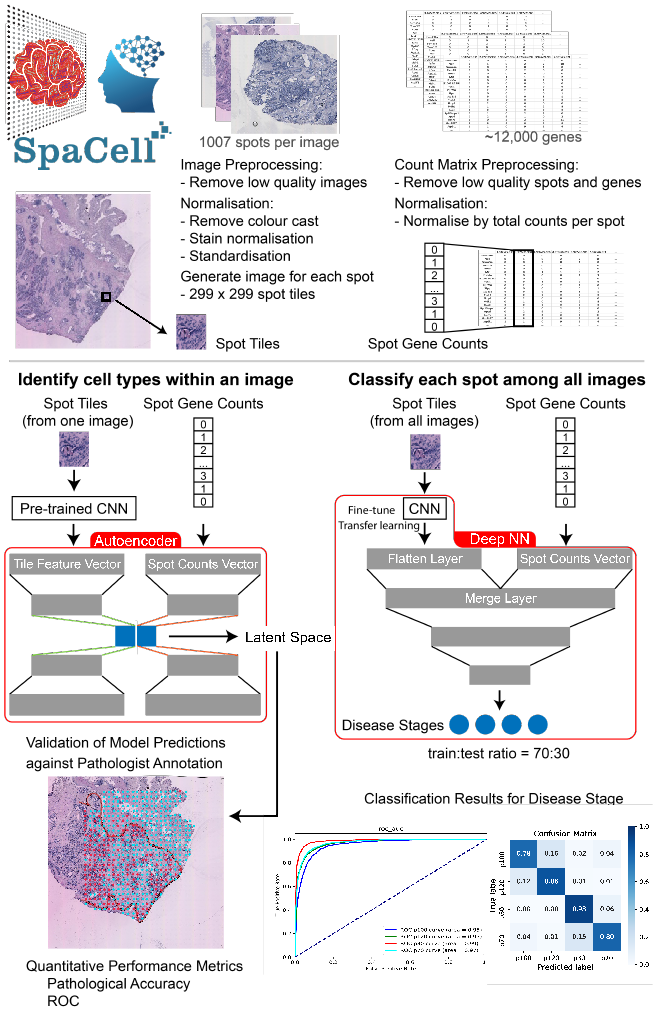
Two main analysis workflows of SpaCell: cell type identification and disease stage classification.

In the cell type classification model (Supp. Methods), SpaCell analyses one high-resolution image and its spatial count matrix. To extract a latent feature vector for each image tile, we fit pre-trained convolutional neural network (CNN) weights from the ResNet50 model to utilize big data in the ImageNet database. For each tile feature vector and its corresponding spot gene counts, we trained two autoencoders (AE) to find two latent spaces of equal dimension, which are then concatenated into one latent vector. For all spots, the latent vectors are then combined to form a latent matrix representative for both image and gene count data, which are then used to perform clustering (default as K-means clustering) to identify cell types. After spot clustering, SpaCell provides visualisation functions to evaluate the model performance. Importantly, to enable quantitative comparison to pathological annotation information, we devised an approach to automatically and accurately detect and map annotation contours from a low-resolution image to a WSI. After mapping, we use digital spot coordinates and the contour-masked regions to compare computational prediction with pathological annotation.

In the disease stage prediction model (Supp. Methods), SpaCell uses hundreds of images and corresponding count matrices. Both tiles and count data are input into a fully connected model, initially as two streams. Each image tile is initiated by a CNN with weights pre-trained on the ImageNet dataset and these weights are trainable together with parameters from the count stream. To increase model generalisation and reduce over-fitting, the following strategies are applied: random sampling of images stratified by labels ensuring unseen test images, five-fold cross-validation, drop-out and L2 penalization. Following model training, users can apply evaluation functions in SpaCell for quantitative analysis of model performance such as test accuracy, ROC curves and confusion matrix.

SpaCell models were tested on a prostate cancer (Berglund *et al.*, 2018) and amyotrophic lateral sclerosis (ALS) (Maniatis *et al.*, 2019) datasets (Supp. Methods), which represent a dataset with few images and high resolution compared to a dataset with more images and lower resolution. By testing more than 40 models, we consistently found that the combination of pixel and gene expression data improved model performance by 8-14% in accuracy, precision, F-score and Area Under the Curve in cell-type models (Supp. Figs 1, 2) and 4% in disease-stage classification models (Supp. Fig 3).

## 3 Implementation

SpaCell has been developed with Python 3.7 as a user-friendly software. Installation and tutorials are described in the SpaCell GitHub page and PyPI repository. Changes in the parameter settings are kept in the config file for reproducibility. SpaCell uses Keras and TensorFlow backend which are portable between platforms and supports Graphics Processing Units (GPUs) distribution to accelerate the training step.

## 4 Conclusion

SpaCell is a pioneering software program implementing deep neural networks for integrating image pixel data and spatial gene expression data for biomedical research. We show that SpaCell can automatically and quantitatively identify cell types and disease stages. We tested over 40 models and consistently found that the integration of both data types increased model performance compared to using one type of data input. Moreover, SpaCell prediction results have higher resolution, specific to thousands of spatial spots, compared to typical pathological annotation with several large regions. We expect that our model can be applied to any type of spatial omics data that have both images and expression values.

## Acknowledgements

We thank Prof. Joakim Lundeberg and Dr. Emelie Berglund for sharing the spatial data and members in Nguyen’s Biomedical Machine Learning Lab for helpful discussion. This work has been supported by the Australian Research Council (ARC DECRA DE190100116), the University of Queensland, and the Genome Innovation Hub.

## Supplementary information

### Supplementary methods

#### Cell Type Clustering

Data preprocessing includes tiling and normalisation to solve two inherent challenges, the small sample size and the technical variation between images. Since the number of images is often small, SpaCell implements a tiling strategy where each spatial spot in Slide-seq data is captured as an image tile with a corresponding column in the count matrix. While most histological images are viewed in isolation without taking into account other images, SpaCell can use spot image tiles from many tissue section images. Images are often different in contrast, colour, and baseline brightness due to various factors such as manufacturers, microscopy settings and slide preparation. The normalisation step in SpaCell removes this variation through colour cast removal and stain normalisation. As histological images are taken against a white background, any images with a coloured background can be assumed to have a colour cast. To remove this colour cast, SpaCell scales the R,G,B channels individually such that the background becomes white. Stain normalisation is implemented in StainTools as described below (Macenko *et al.*, 2009):

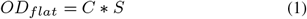

where *OD* is the optical density image transformed from a H&E stained image in RGB format, *S* is a stain two colour matrix, and *C* is a pixel concentration matrix that is used to normalise a target image using the *S* matrix of that image. By default, the stain matrix is estimated by Vahadane *et al.* method (Vahadane *et al.*, 2016), or alternatively by Macenko *et al.* method (Macenko *et al.*, 2009). We found at least 3% improvement in accuracy for models with the normalisation step.

In the cell type identification model, SpaCell uses a pre-trained convolutional neural network (CNN), ResNet50, which makes use of network weights trained from the ImageNet database. SpaCell applies the ResNet50 model (He *et al.*, 2015) to each spot image to find a latent variable vector representing informative features in each spot image. Due to the unbalanced number of features in the gene counts (13,000 genes) and tile feature vector (2048 features) for each spot, SpaCell has function to select 2048 top variable genes to equalise the dimensions of the spot gene counts and spot tile feature vector to 2048. SpaCell also performs min-max scaling method to scale spot gene counts and spot tile feature vector to range of 0 to 1 to minimize bias derived from variation in data ranges between image and count data. SpaCell implements two separate autoencoders for gene count and tile feature vector data to generate two latent spaces with the same dimensions. Those two latent spaces are able to output key features representative of both the original spot image and the spot gene expression, therefore, are concatenated as input for downstream k-means clustering (Fig. 1). SpaCell implements three loss functions such as the Mean Square Error (MSE) (Equation (2)), Kullback-Leibler (KL) (Equation (3)) and Binary Cross Entropy (BCE) (Equation (4)) to measure the cost of the original input and the constructed output. Loss is minimised by the Adam (Kingma and Ba, 2014) optimiser during the training step (epoch).

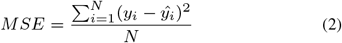

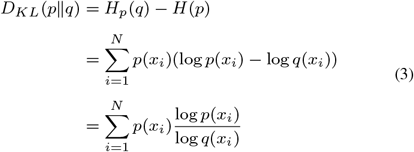

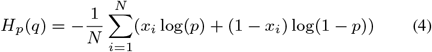

where *i* is the index for spot *i*. *X*_*i*_ represents 2048 feature vectors from ResNet50 model or a vector of counts for the top 2048 most variable genes measured for *Spot_i_*. *p* and *q* denote the probability distributions for input and constructed output of all *N* spots.

To quantify performance of the cell type identification model, SpaCell implements a validation tool that utilises pathologist annotations. Pathologist annotations are often low-resolution so SpaCell registers this annotation image to the whole slide image. To achieve this, SpaCell uses a sliding window approach to find the best location and scale for the annotation image, as indicated by a maximum normalised correlation coefficient (Yoo and Han, 2009). In combination with a user-specified annotation colour, SpaCell extracts the annotation contours. These annotations may be open or closed contours. SpaCell preprocesses open contours with a convex hull approach (Barber *et al.*, 1996) to close the contours. Closed contours are filled in to create an annotation mask. By referencing spot coordinates against this pathological annotation mask, SpaCell generates a pathologist label for each spot. SpaCell compares the labels predicted by the cell type identification model to the pathologist labels to generate performance metrics such as accuracy, F-score and ROC curves.

Clustering method were tested on a prostate cancer dataset (Berglund *et al.*, 2018) containing 12 tissue slides from one patient but taken from different prostate locations. Two slides with pathologist annotation were used to test the model performance; P3.3 which represents cancer and non-cancer regions with an open annotation contour and P4.4 which represents inflamed stromal and non-inflammed stromal regions with a closed annotation contour.

#### Disease Stage Classification

In the disease stage prediction model, SpaCell uses a two-input deep neural framework to integrate spot image data and spot gene count data (Fig. 1). Spot images feed into an ResNet50 followed by a hidden layer for image feature input and spot gene counts feed into a hidden layer for gene expression input. A merged layer connects these two hidden layers and is followed by a fully connected neural network classifier. This architecture enables the model to learn effective information from both spot image data and spot count data that is relevant to the disease stage. This model uses a Softmax activation function in the last layer to calculate the probability over the *C* classes for each input *Z*, defined as:

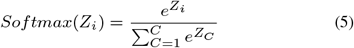

As this model is designed for multi-class classification problem, a Categorical Cross-Entropy Loss (CCE) (Equation (6)) is implemented where *t_i_* and *Softmax*(*Z_i_*) denote the ground truth and predicted score for class *C_i_*.

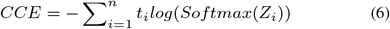

Dropout layer and l2 regularization for dense layer are used to avoid overfitting. To improve the robustness of the model and assess the risk of overfitting, SpaCell implements stratified 5 fold cross-validation on the training dataset.

Classification method were tested on ALS dataset (Maniatis *et al.*, 2019) consisting of 348 Spinal cord spatial transcriptomics tissue slides in the raw dataset, of which 146 were classified into different ALS disease stages. After removing low quality spots, 31771 spots within 143 WSIs were kept for classification model. Each WSI was labeled as one of the four disease progression categories, including p30, p70, p100 and p120 representing pre-symptomatic, onset, symptomatic and end-stage respectively. In total, 9267 tiles from 46 WSIs with p30 labelled, 6370 tiles from 33 WSIs with p70 labelled, 6337 tiles from 27 WSIs with p100 labelled and 9797 tiles from 37 WSIs with p120 labelled were used in classification model. 143 ALS mice were randomly split to 100 mice for training and the remaining 43 mice as unseen data for testing such that each disease stage was represented in similar proportions in the test set. Five fold cross validation was applied for the training dataset. After tilling without overlapping, 22179 tiles from 100 mice were used for model training and 9592 tiles from 43 unseen mouse tissue images were used for testing the model performance.

### Supplementary results

#### Compare models for cell type classification

For cell type clustering model, we assessed SpaCell’s performance in two applications different in biological contexts. In one case, we distinguished cancer cells from non-cancer cells (image P4.2) and in another case we identified stromal cells from the whole tissue (image P3.3), (Supp. Fig 1). Two images were selected because they had pathological annotation, which can be used as a reference for assessing spot predicted values. We successfully mapped the contours from low-resolution pathological annotation images, available as pdf files in the original paper (Berglund *et al.*, 2018), to the original images that are 1000 times larger. The mapping enables us to assess the model performance by accuracy, precision, F-score, and ROC curve. In both cases, the combined model shows higher performance than using one data type alone, with up to 25% in precision, 14% in accuracy and 38% for F-score. Furthermore, we compared 36 models, with different options for data inputs (combined with two latent spaces, combined with one latent space, single gene-count, single tiled image), data-prepossessing (scaled, top variable genes, and no-scaled), and loss functions (BCE, KL, and MSE) (Supp. Fig 2). The four heatmap blocks demonstrate the comprehensive comparisons and the superior performance of the combined pixel and gene-count models, with the performance ranked in descending order as: combined with two latent spaces, combined with one latent space, single gene-count model, and single tiled images (Supp. Fig 2). We also found the optimal architecture for the cell type classification model with two latent spaces, BCE loss, and scaled preprocessing. With this optimal model, the performance of the combined image and gene-count data is 8-14 % higher in accuracy, precision, F-score and Area Under the Curve than the models with only one data type (Supp. Fig 2).

#### Compare models for disease stage classification

For multiclass classification of the four ALS disease stages, we implemented a stringent design to create a test set completely unseen from the training dataset at both the tile and image levels and performed cross-validation to assess overfitting and model robustness. The design allowed us to assess model performance based on ground-truth labels from known phenotype for each of the above 31,000 tiles representing 143 mice and four disease states. Supp Fig. 3 shows higher performance for the combined model especially for distinguishing the to very similar class, presymtomatic (p30) and onset (p70). The confusion matrix in the supplementary Figure 3 B show that the gene count only model was unable to separate these two classes (P30 and P70). The combined model also performed markedly better compared to the model using image only input.

### Implementation steps

~~~
# Step 1. Installation
git clone https://github.com/BiomedicalMachineLearning/Spacell.git
conda create -y -name spacell python==3.7
conda install -y -name spacell -c conda-forge --file requirements.txt
conda activate spacell
~~~

~~~
# Step 2. Setup configurations in config.py
# Path to metadata which contains at least sample name column and
# corresponding label column
META_PATH = ’‥/dataset/metadata/mouse_sample_names_sra.tsv’
~~~

~~~
# Path to spatial transcriptomics imaging data
IMG_PATH = ’‥/dataset/image/’
~~~

~~~
# Path to spatial transcriptomics gene counts data
CM_PATH = ’‥/dataset/cm/’
~~~

~~~
# Alignment transform matrix
# If ST imaging data were aligned, leave it to None.
# Otherwise, give the path to affine transformation
# matrix generated by st_spot_detector
ATM_PATH = None
~~~

~~~
# Path to folder that save the tiles
TILE_PATH = ’‥/dataset/tile/’
~~~

~~~
# Path to save intermediate output and final result
DATASET_PATH = ’‥/dataset/’
~~~

~~~
# Path to an image which will be used as a template
# for stain normalization
TEMPLATE_IMG = ’‥/dataset/image/CN94_D2_HE.jpg’
~~~

~~~
# Tile size (DO NOT CHANGE)
SIZE = 299, 299
~~~

~~~
# Color channel (RGB)
N_CHANNEL = 3
~~~

~~~
# Image stain normalization method
NORM_METHOD = ’vahadane’
~~~

~~~
# Threshold for removing low abundant gene, genes
# expressed in less than THRESHOLD_GENE of total
# number spots will be removed
THRESHOLD_GENE = 0.01
~~~

~~~
# Threshold for removing low quality spots, spots
# with less than THRESHOLD_SPOT of total genes
# expressed will be removed
THRESHOLD_SPOT = 0.01
~~~

~~~
# Minimum gene count value for counting whether
# expressed or not
MIN_EXP = 1
~~~

~~~
# Specify column name of sample name column,
# label column and condition column (used
# for subset if provided otherwise leave it to None)
# in metadata file
SAMPLE_COLUMN = ’sample_name’
LABEL_COLUMN = ’age’
CONDITION_COLUMN = ’breed’
~~~

~~~
# Subset dataset that all samples have certain
# CONDITION in CONDITION_COLUMN
CONDITION = ’B6SJLSOD1-G93A’
ADDITIONAL_COLUMN = 2 if CONDITION_COLUMN else 1
~~~

~~~
# Set random seed for reproducibility
seed = 37
~~~

~~~
# Color map for spots color in final clustering plot
color_map = [’#ff8aff’, ’#6fc23f’, ’#af63ff’,
’#eaed00’, ’#f02449’, ’#00dbeb’, ’#d19158’,
’#9eaada’, ’#89af7c’, ’#514036’]
~~~

~~~
# Options for running different models:
# “combine” : uses both image and gene count data
# “gene_only” : takes gene count data only as input
# “tile_only” : takes images data only as input
model = [“combine”, “gene_only”, “tile_only”]
~~~

~~~
# Number of tiles that will be propagated through
# the model at each step
batch_size = 32
~~~

~~~
# Number of times that all training dataset will be
# passed forward and backward through model
epochs = 1
~~~

~~~
# The ratio for splitting training and test datasets
train_ratio = 0.5
~~~

~~~
# Number of categories in label
n_classes = 4
~~~

~~~
# Option for stratified K-fold
# True : run stratified K-fold cross validation
# False : run model without cross validation
~~~

~~~
cross_validation = False
~~~

~~~
# Number of splits for cross validation
k_fold = 2
~~~

~~~
# Step 3. Image Preprocessing
python image_normalization.py
~~~

~~~
# Step 4. Count Matrix PreProcessing
python count_matrix_normalization.py
~~~

~~~
# Step 5. Generate paired image and gene count
# training dataset
python dataset_management.py
~~~

~~~
# Step 6. Classification
python spacell_classification.py
~~~

~~~
# Step 7. Clustering
python spacell_clustering.py -i /path/to/one/image.jpg -l
/path/to/iamge/tiles/ -c /path/to/count/matrix/
-e 100 -k 2 -o /path/to/output/
~~~

~~~
# -e is number of training epochs
# -k is number of expected clusters
~~~

~~~
# Step 8. Clustering Validation and Quantification
python spacell_validation.py -d /path/to/data
-a annotation.png -w wsi.jpeg
-m affine_tranformation_matrix.txt
~~~

~~~
# -o output_folder
# -k clustering_predictions.tsv
# -c annotation_colour_range
# -c is annotation colour range thresholds
# -blue_low green_low red_low blue_upper green_upper red_low
# -t indicates that annotations are not closed paths,
# so spacell with try to close the paths
# -f downscale factor if the input whole slide image has
# already been downscaled
# -s spot size, optional, usually set automatically
~~~

## Supplementary materials

**Supplementary Figure 1.**
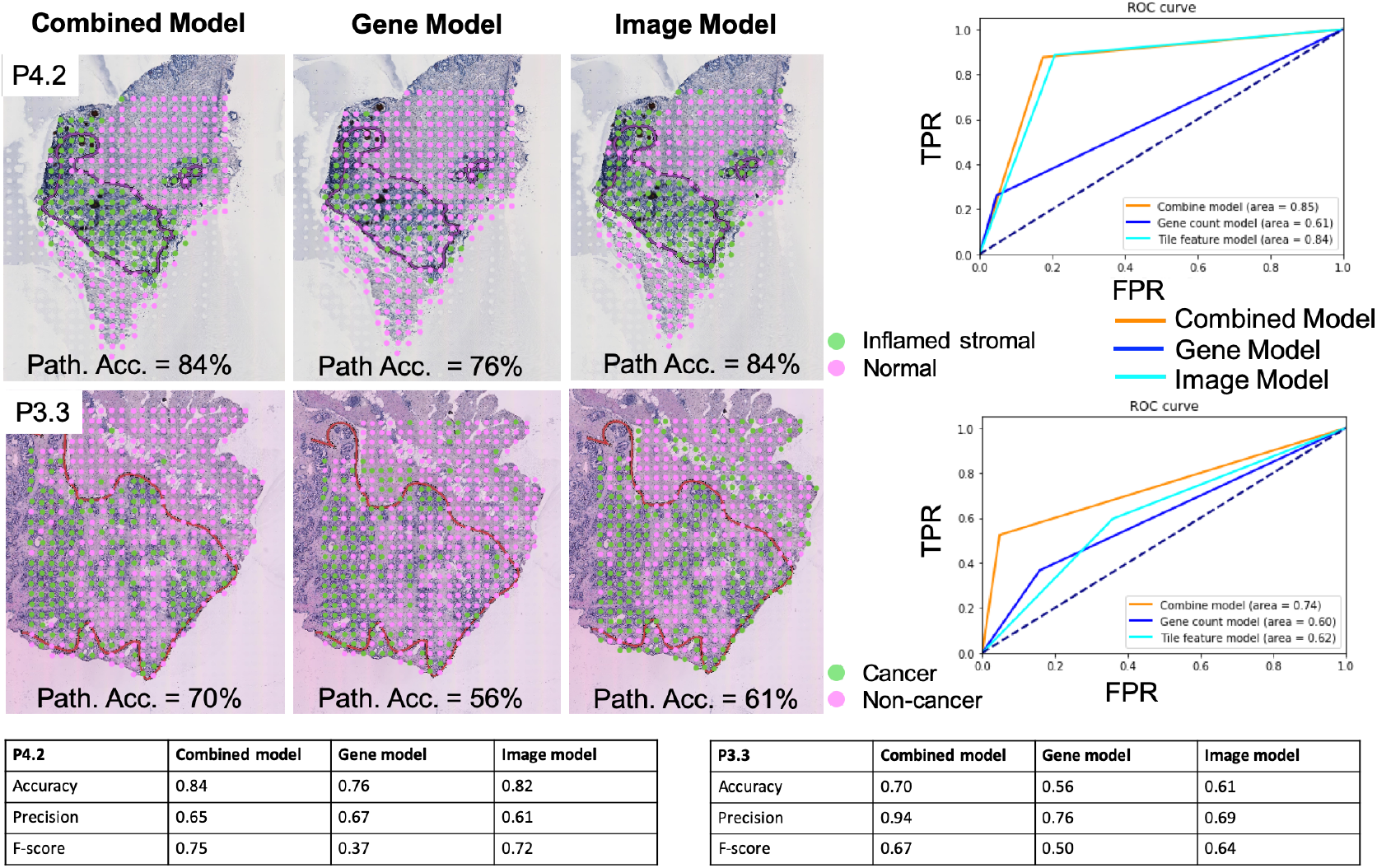
Cancer and inflamed cell type classification. Comparisons between three models that used either only gene count data or image data or the combination of both as the model training dataset. P4.2, inflamed stromal cell. P3.3, cancer cell types. The red and purple contours denote pathological annotation. All three models implemented same AE architecture with 100 training epochs where losses were calculated by BCE. Top 2048 variable genes were selected in gene model and combined model to balance the weight of gene expression data and tile feature data.

**Supplementary Figure 2.**
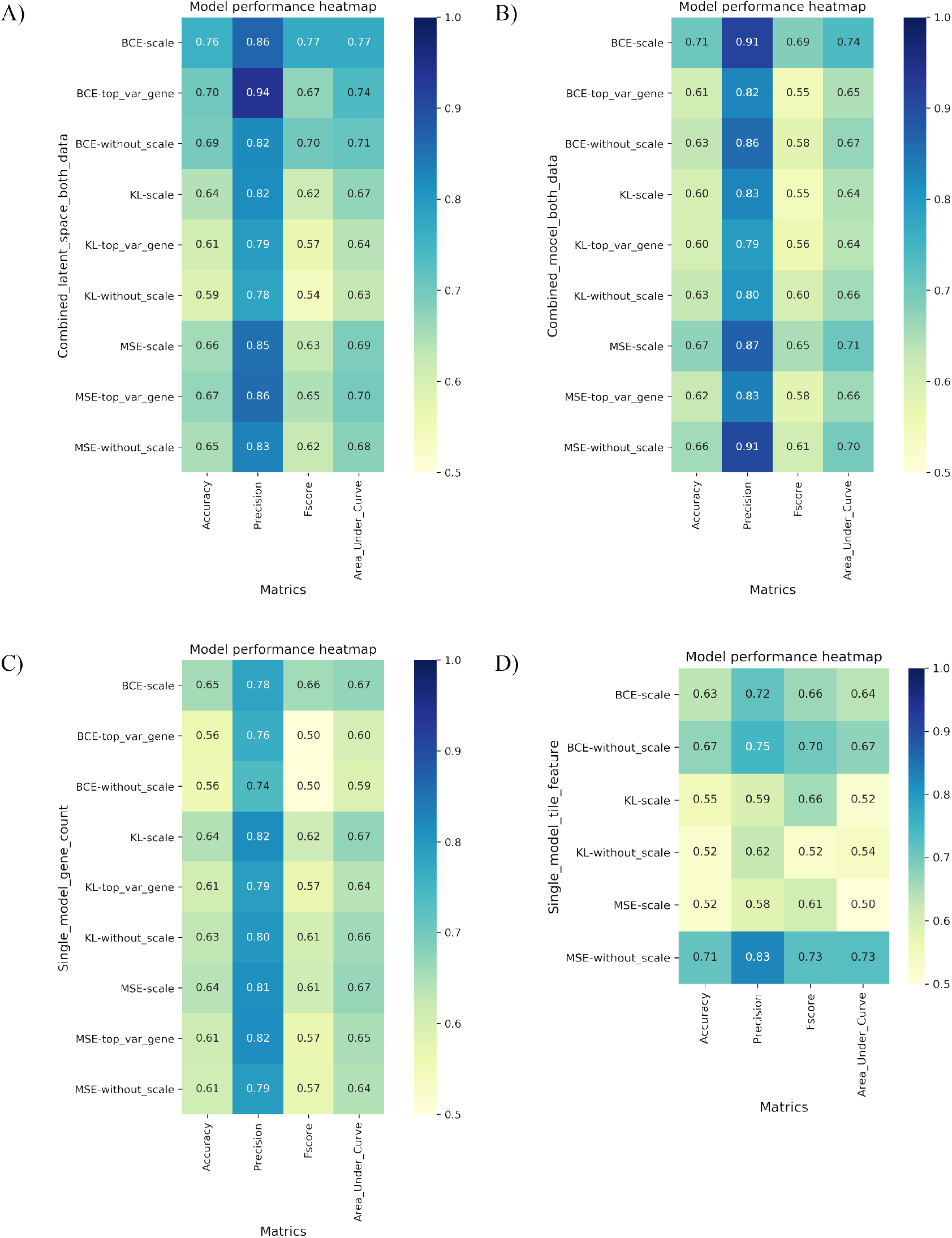
Comparing cell type classification models. Four data input categories are compared. A) is model using the combination of gene counts and images with two separate autoencoder streams, followed by the concatanation of the two latent spaces. B) is model using the combination of gene counts and images with a single autoencoder used for both pixel and gene count. C) and D) are gene count only model and image only model, respectively.

**Supplementary Figure 3.**
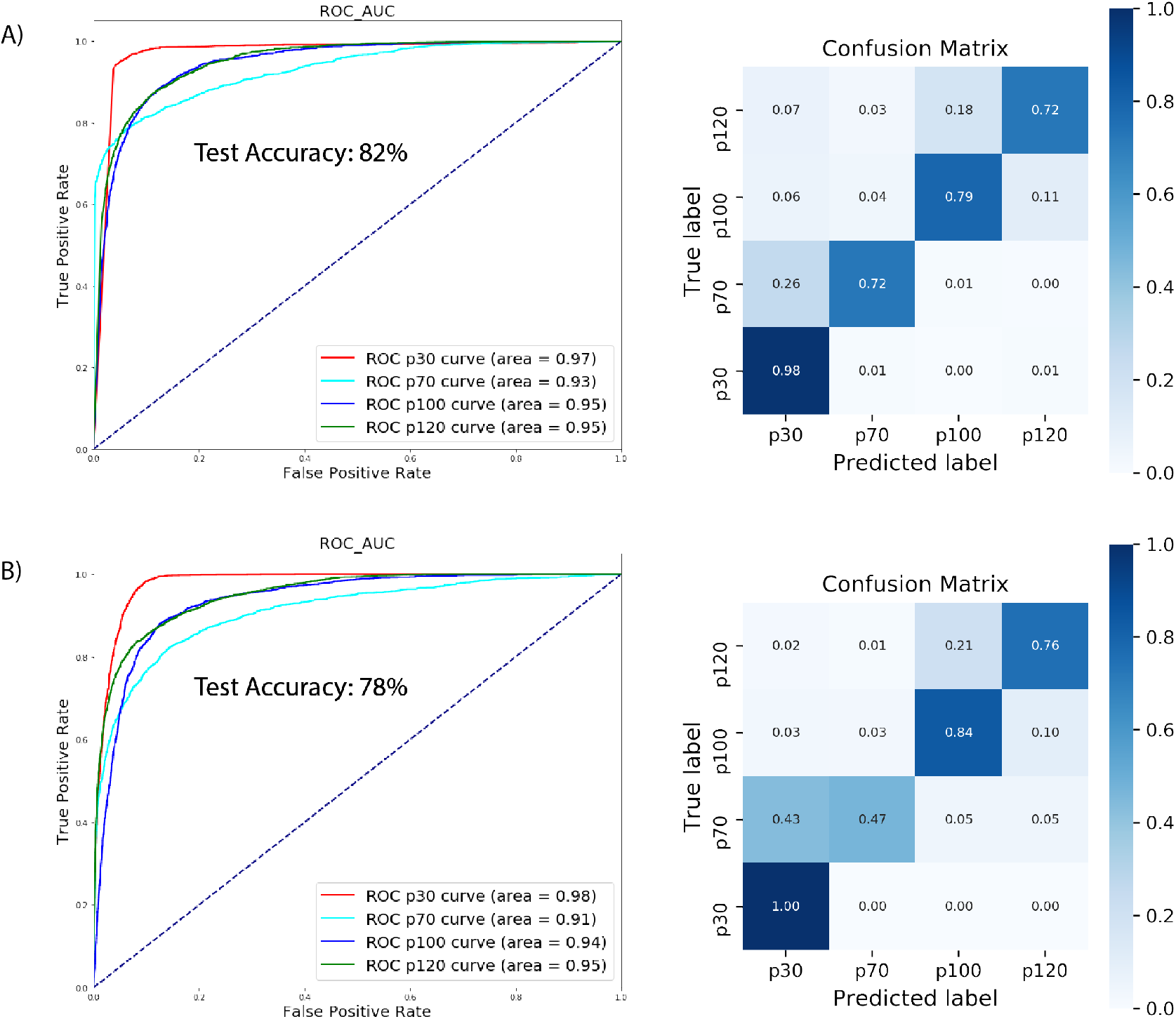
Disease stage classification model. Three models are compared. A) and B) are the model using the combination of gene counts and images or the gene count only, respectively.

